# Structures of potent and convergent neutralizing antibodies bound to the SARS-CoV-2 spike unveil a unique epitope responsible for exceptional potency

**DOI:** 10.1101/2020.07.09.195263

**Authors:** Shuo Du, Yunlong Cao, Qinyu Zhu, Guopeng Wang, Xiaoxia Du, Runsheng He, Hua Xu, Yinghui Zheng, Bo Wang, Yali Bai, Chenggong Ji, Ayijiang Yisimayi, Qisheng Wang, Ning Gao, X. Sunney Xie, Xiao-dong Su, Junyu Xiao

## Abstract

Understanding the mechanism of neutralizing antibodies (NAbs) against SARS-CoV-2 is critical for effective vaccines and therapeutics development. We recently reported an exceptionally potent NAb, BD-368-2, and revealed the existence of *VH3-53/VH3-66* convergent NAbs in COVID-19. Here we report the 3.5-Å cryo-EM structure of BD-368-2’s Fabs in complex with a mutation-induced prefusion-state-stabilized spike trimer. Unlike *VH3-53/VH3-66* NAbs, BD-368-2 fully blocks ACE2 binding by occupying all three receptor-binding domains (RBDs) simultaneously, regardless of their “up” and “down” positions. BD-368-2 also triggers fusogenic-like structural rearrangements of the spike trimer, which could impede viral entry. Moreover, BD-368-2 completely avoids the common epitope of *VH3-53/VH3-66* NAbs, evidenced by multiple crystal structures of their Fabs in tripartite complexes with RBD, suggesting a new way of pairing potent NAbs to prevent neutralization escape. Together, these results rationalize a unique epitope that leads to exceptional neutralization potency, and provide guidance for NAb therapeutics and vaccine designs against SARS-CoV-2.

## Introduction

Coronavirus disease 2019 (COVID-19), caused by the severe acute respiratory syndrome coronavirus 2 (SARS-CoV-2), has become a global pandemic (Callaway et al., 2020). An important structural protein of SARS-CoV-2 is the Spike (S) protein, which recognizes human angiotensin-converting enzyme 2 (ACE2) to mediate the fusion between viral and host cell membranes (Hoffmann et al., 2020; Walls et al., 2020). The S protein could be divided into two regions, S1 and S2. S1 contains the N-terminal domain (NTD), which likely contributes to maintaining the prefusion state of the S protein, and the receptor-binding domain (RBD), which is responsible for interacting with ACE2 (Lan et al., 2020; Shang et al., 2020; Wang et al., 2020b; Xu et al., 2020; Yan et al., 2020; Zhou et al., 2020b). Binding of ACE2 to RBD induces a conformational change in the S protein, leading to the exposure of the membrane fusion peptide in S2 that subsequently functions in the membrane fusion process. Structural analyses of the S trimer reveals that RBDs could adopt different “up” and “down” conformations (Ke et al., 2020; Walls et al., 2020; Wrapp et al., 2020), which has important implications in both receptor binding and immune recognition.

Neutralizing antibodies are important therapies for COVID-19. SARS-CoV-2 neutralizing antibodies (NAbs) targeting the RBD, as well as the NTD, were reported extensively (Barnes et al., 2020; Brouwer et al., 2020; Cao et al., 2020; Chi et al., 2020; Hansen et al., 2020; Ju et al., 2020; Liu et al., 2020; Pinto et al., 2020; Robbiani et al., 2020; Rogers et al., 2020; Seydoux et al., 2020; Shi et al., 2020; Wang et al., 2020a; Wec et al., 2020; Wu et al., 2020; Zhou et al., 2020a). Recently, we identified a series of potent neutralizing antibodies from convalescent patients using high-throughput single-cell RNA sequencing (Cao et al., 2020). The most potent one, BD-368-2, exhibited high therapeutic and prophylactic efficacy in hACE2-transgenic mice infected by SARS-CoV-2 (Bao et al., 2020). We also revealed the wide existence of a convergent, public, or stereotypic antibody response to SARS-CoV-2 by *VH3-53/VH3-66* derived NAbs, which was confirmed and highlighted in several recent studies (Barnes et al., 2020; Hansen et al., 2020; Kim et al., 2020; Robbiani et al., 2020; Rogers et al., 2020; Yuan et al., 2020a). The *VH3-53/VH3-66* convergent NAbs share highly conserved VDJ sequences and could be found in different individuals in distinct populations, similar to what has been observed in HIV, Influenza, and Hepatitis C viruses (Ekiert et al., 2009; Gorny et al., 2009; Marasca et al., 2001).

Since BD-368-2 and several *VH3-53/VH3-66* NAbs, such as CB6 (Shi et al., 2020), are now undergoing clinical evaluations, it is critical to understand their detailed interactions with the S protein and the molecular mechanisms behind their high neutralizing potency. On the other hand, gaining further insights into the interactions between these NAbs and the S protein could also facilitate the design of S protein variants that are stabilized at particular conformations to be used as vaccines, as demonstrated by the success of structure-based vaccine design for the respiratory syncytial virus (Crank et al., 2019; Graham et al., 2019). Here we conducted a systematic structural analysis of the antibody response to the S protein. We obtained a large collection of *VH3-53/VH3-66* and *JH4/JH6* derived antibodies and showed that a high proportion of these antibodies are potent SARS-CoV-2 NAbs. Multiple high-resolution crystal structures of the RBD in complex with the Fabs of these NAbs revealed that the convergent NAbs all share a highly conserved epitope that is only accessible in the “up” RBDs.

We also investigated BD-368-2’s neutralizing mechanism and its molecular interaction with the S protein. With the help of a mutation-induced prefusion-state-stabilized S trimer, we determined the 3.5-Å cryo-EM structure of BD-368-2 Fabs in complex with the S trimer and showed that BD-368-2 fully blocks ACE2 binding by occupying all three RBDs simultaneously regardless of their “up” and “down” positions. Furthermore, the tripartite crystal structures of RBD in complex with the Fabs of both BD-368-2 and *VH3-53/VH3-66* NAbs demonstrated that their epitopes are entirely non-overlapping. These results rationalize the potent neutralizing mechanism of BD-368-2 and suggest the ideal therapeutic application of BD-368-2 in pair with the *VH3-53/VH3-66* NAbs to prevent neutralization escape caused by mutation of the S protein.

## Results

### *VH3-53/VH3-66* derived convergent antibodies exhibit high neutralizing potency

Previously, we reported a phenomenon of a stereotypic/convergent antibody response to SARS-CoV-2. Those convergent antibodies were derived from the *VH3-53/VH3-66* family and showed a high likelihood of being SARS-CoV-2 NAbs. Multiple groups have reported similar NAbs, such as CB6, demonstrating their wide existence in distinct populations. More importantly, the NAbs generated by different individuals not only share highly similar V_H_ gene segments, but also exhibit conserved sequences for both J_H_ and V_L_ gene segments. To gain more insights into the convergent NAbs, we synthesized 28 additional *VH3-53/VH3-66* derived antibodies from the RBD-enriched high-throughput single-cell sequencing library (Cao et al., 2020). All antibodies were selected based on their variable (V), diversity (D), and joining (J) combinations, which contain a *VH3-53/VH3-66* V_H_ gene, a *JH4/JH6* J_H_ gene, and a *VK1-9/VK1D-33/VK1D-39/VK3-20* V_L_ gene. Combined with the previously reported antibodies, such as BD-236, a total of 45 *VH3-53/VH3-66* derived antibodies were collected (Table S1). Verified by a pseudovirus-based neutralization assay, we found that nearly two-thirds of the *VH3-53/VH3-66* antibodies displayed neutralizing abilities against SARS-CoV-2 (Figure 1A). Also, seven new potent NAbs were discovered, all showing an IC_50_ below 20 ng/mL (Figure 1B, Figure S1) and a high binding affinity for RBD (Figure S2). Together, our collection of *VH3-53/VH3-66* derived convergent NAbs showed clear evidence for the existence of a strong, recurrent antibody response to SARS-CoV-2.

**Figure 1.**
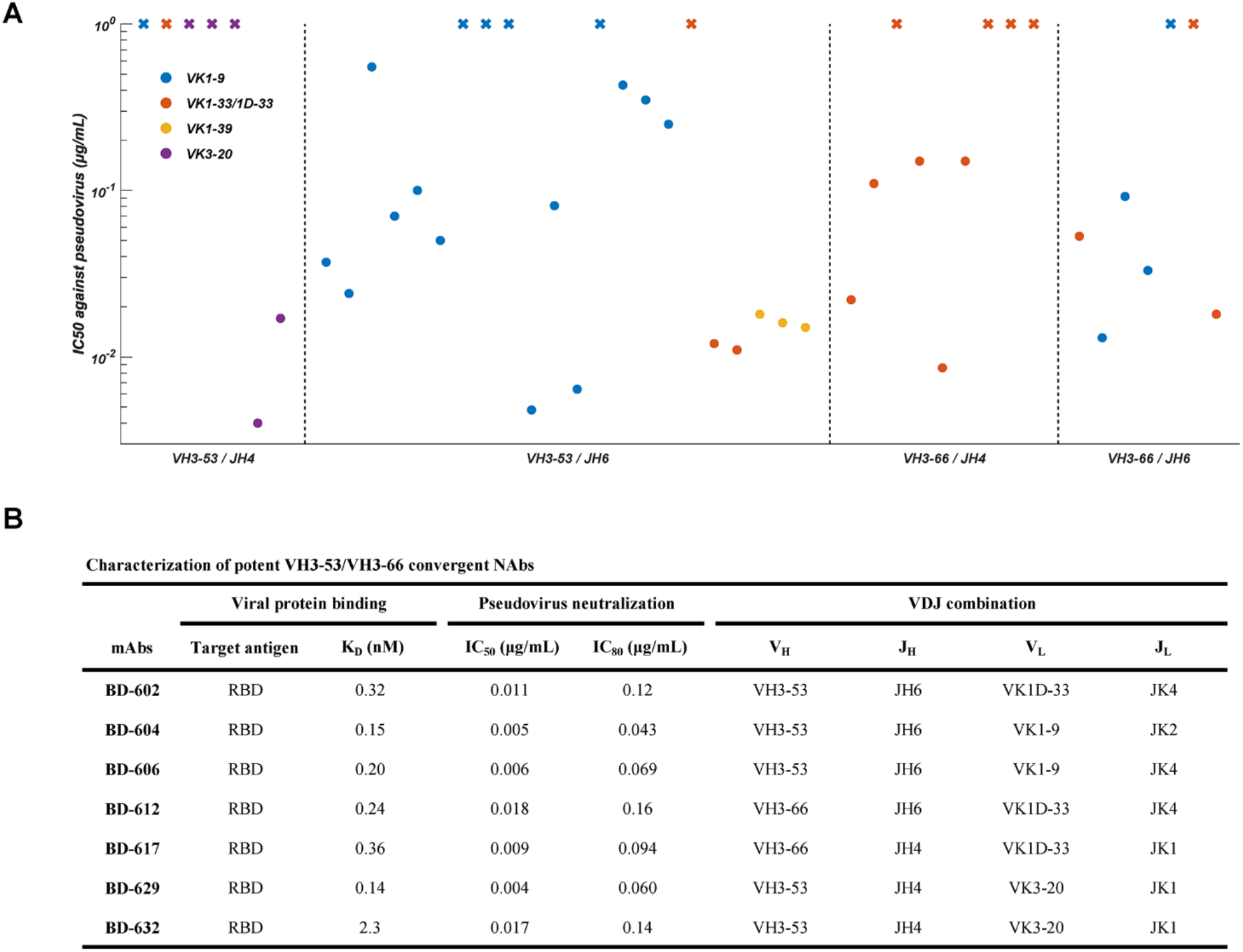
Binding affinity and neutralizing abilities of *VH3-53/VH3-66* derived antibodies. (A) The distribution of IC_50_ against SARS-CoV-2 pseudovirus for *VH3-53/VH3-66* derived antibodies revealed by high-throughput single-cell sequencing. Data for each antibody were obtained from a representative neutralization experiment, which contains three replicates. IC_50_ was calculated by using a four-parameter logistic curve-fitting and represented as mean. Each antibody’s heavy chain V-J gene is indicated on the x-axis, where the light chain V gene is indicated by different colors, as shown in the legend. A cross mark indicates that the antibody’s IC_50_ is higher than 1 μg/mL. The detailed characteristics of the antibodies shown here are listed in Table S1. (B) Characteristics of the potent *VH3-53/VH3-66* convergent NAbs selected based on VDJ sequences. K_D_ targeting RBD was measured by using surface plasmon resonance (SPR) with a 1:1 binding model. See also Figure S1 and Figure S2.

### Crystal structures of *VH3-53/VH3-66* antibodies in complex with RBD

To characterize the molecular interactions between the *VH3-53/VH3-66* NAbs and RBD, we determined the crystal structures of RBD in complex with the antigen-binding fragment (Fabs) of BD-236, BD-604, and BD-629 at 2.4 Å, 3.2 Å, and 2.7 Å, respectively (Table S2). These Fabs bind to the RBD with almost identical poses and impose very similar footprints on RBD (Figure 2). Furthermore, they bind to the RBD in similar fashions as B38 (Figure 2D), a *VH3-53* antibody (Wu et al., 2020); and CB6 (Figure 2E), a *VH3-66* antibody (Shi et al., 2020). The structures of several other *VH3-53/VH3-66* mAbs have also been recently reported, including C105 (Barnes et al., 2020), CC12.1 and CC12.3 (Yuan et al., 2020a), and CV30 (Hurlburt et al., 2020). They all appear to interact with RBD in very similar manners (Figure S3). The abundant presence of the *VH3-53/VH3-66* NAbs in patients, and the fact that these antibodies target a commonly shared epitope on the RBD, suggesting that this region of the S protein is highly effective in eliciting immune responses.

**Figure 2.**
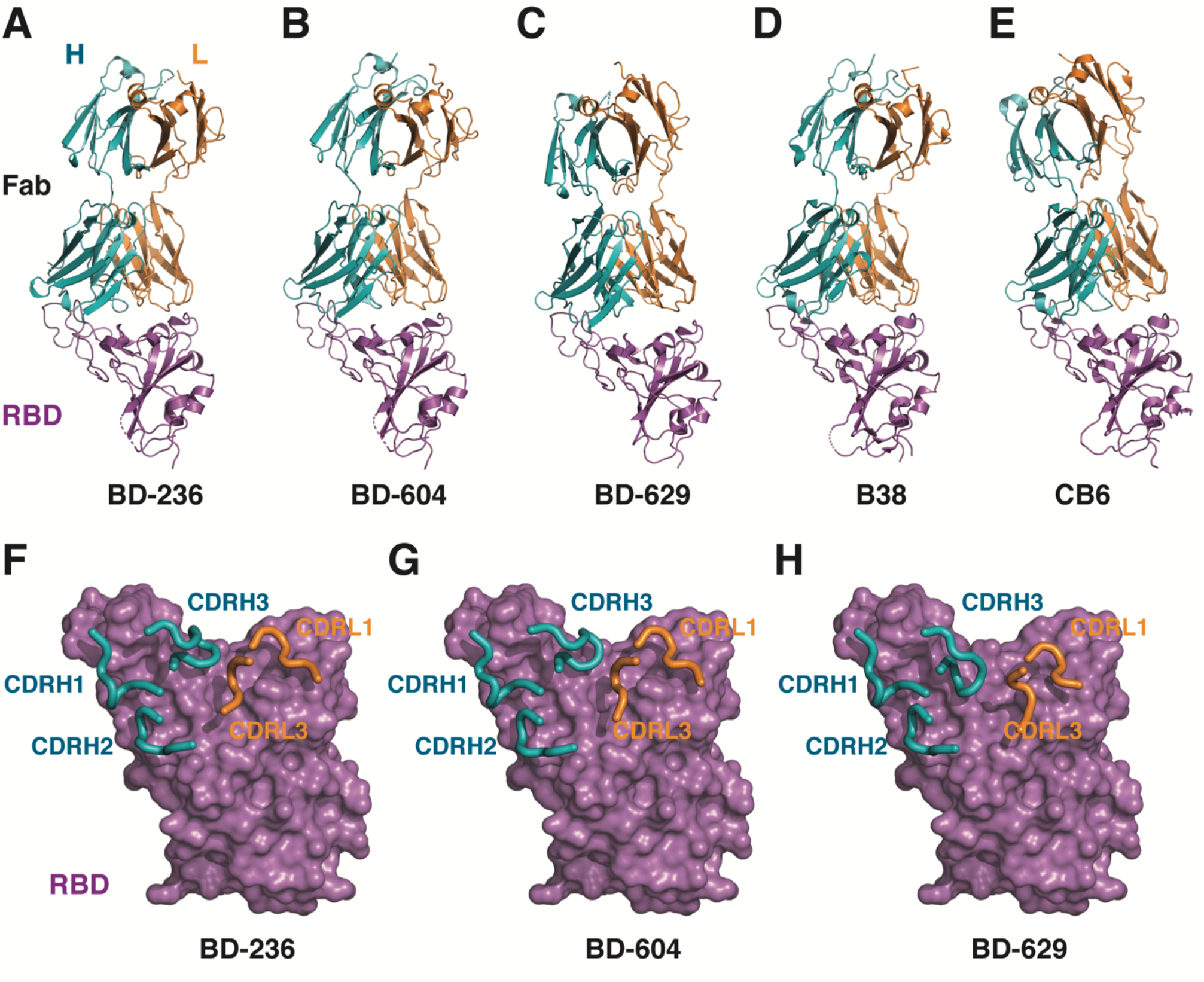
The *VH3-53/VH3-66* antibodies bind to RBD in a similar manner. (A) The crystal structure of BD-236 Fab in complex with RBD. The heavy chain (H) and light chain (L) of BD-236 Fab are shown in teal and orange, respectively. The RBD is shown in magenta. Disordered regions are depicted as dashed lines. (B) The crystal structure of BD-604 Fab in complex with RBD. (C) The crystal structure of BD-629 Fab in complex with RBD. (D) The crystal structure of B38 Fab in complex with RBD (PDB ID: 7BZ5). (E) The crystal structure of CB6 Fab in complex with RBD (PDB ID: 7C01). (F) Three heavy chain CDRs (CDRH1-3) and two light chain CDRs (CDRL1, CDRL3) in BD-236 Fab mediate the interaction with RBD. The CDRs are highlighted using thicker ribbons. RBD is shown in a surface view. (G) Interactions between BD-604 Fab and RBD. (H) Interactions between BD-629 Fab and RBD.

BD-236, BD-604, and BD-629 all interact with the RBD using their three heavy chain CDRs (CDRH1-3) and two light chain CDRs (CDRL1, CDRL3) (Figure 2F-H). Differences in these regions, especially the heavy chain CDRs, account for their different binding affinities to RBD. For example, BD-236 and BD-604 are highly similar to each other. Their immunoglobulin heavy chain variable (V), and joining (J) regions are both encoded by germline genes *IGHV3-53* and *IGHJ6*. Their light chains are derived from the same V region (*IGKV1-9*) as well, and differ only in the J regions (*IGKJ4* for BD-236, whereas *IGKJ2* for BD-604). Only four and two amino acids are different in their respective CDRHs and CDRLs that are involved in interacting with RBD. Nevertheless, BD-604 binds to RBD with a Kd of 0.15 nM, ~19 fold higher than BD-236, which displays a Kd of 2.8 nM (Table S1). BD-604 is also more potent against the SARS-CoV-2 pseudovirus, with an IC_50_ value of 5 ng/ml, compared to 37 ng/ml for BD-236. Detailed structural analyses reveal that two critical aromatic residues in the CDRH2 and CDRH3 of BD-604 likely contribute to its higher affinity interaction with RBD (Figure S4A). First, Phe58 in CDRH2 replaces Asp58 in BD-236, leading to a better packing with Tyr52 and also van der Waals interactions with Thr415^S^-Gly416^S^ (superscript S indicates the amino acid of the S protein) (Figure S4B). Second, Tyr102 in CDRH3 substitutes for Ala102 in BD-236, and it interacts with Gln493^S^ via a hydrogen bond (Figure S4C). In contrast, even though the heavy chain and light chain genes encoding BD-629 are more different when compared to BD-604 except for *IGHV3-53*, its RBD binding affinity and neutralization ability are similar to BD-604 (Figure 1B, Table S1). Aromatic residues corresponding to Phe58 and Tyr102 in BD-604 are both present in BD-629. In CDRH2, Tyr58 not only mediates packing interactions as described above for Phe58 in BD-604 but also forms hydrogen bonds with Thr415^S^ (Figure S4B). In CDRH3, Tyr102 is present, together with two additional tyrosine residues, Tyr99 and Tyr103 (Figure S4A). Tyr102 is slightly pushed away by Tyr103 to form a hydrogen bond with Tyr453^S^; whereas Tyr99 and Tyr103 mediate packing interactions with Phe456^S^ and Tyr489^S^ (Figure S4C). As a result of these additional interactions, the binding between BD-629 and RBD is more dominated by the heavy chain when compared to BD-604. The VH and VL domains of BD-629 bury 809 and 198 Å^2^ surface areas on RBD, respectively, compared to 754 and 367 Å^2^ for BD-604.

### *VH3-53/VH3-66* antibodies engage the RBDs in the “up” conformation

Importantly, once these antibodies engage the RBD, they would all block ACE2 from binding (Figure 3A), explaining their potent neutralizing activities. When their interaction with RBD is considered in the context of the S trimer, it is evident that these antibodies could only engage the RBDs in the “up” conformation (Figure 3B). Once an RBD is in the “down” position, the attachment of these antibodies would be sterically hindered by an adjacent protomer (Figure 3C). The S trimer has a dynamic nature and can exist in multiple conformations, with the predominant conformations being either close, which has all three RBDs “down”, or partially open, which has one RBD “up” (Ke et al., 2020; Walls et al., 2020; Wrapp et al., 2020). Both of these states would create some conformation barriers for the *VH3-53/VH3-66* NAbs to engage all three RBDs simultaneously, since they would have to seize the stochastic opening moment of the S trimer to snatch the “down” RBDs.

**Figure 3.**
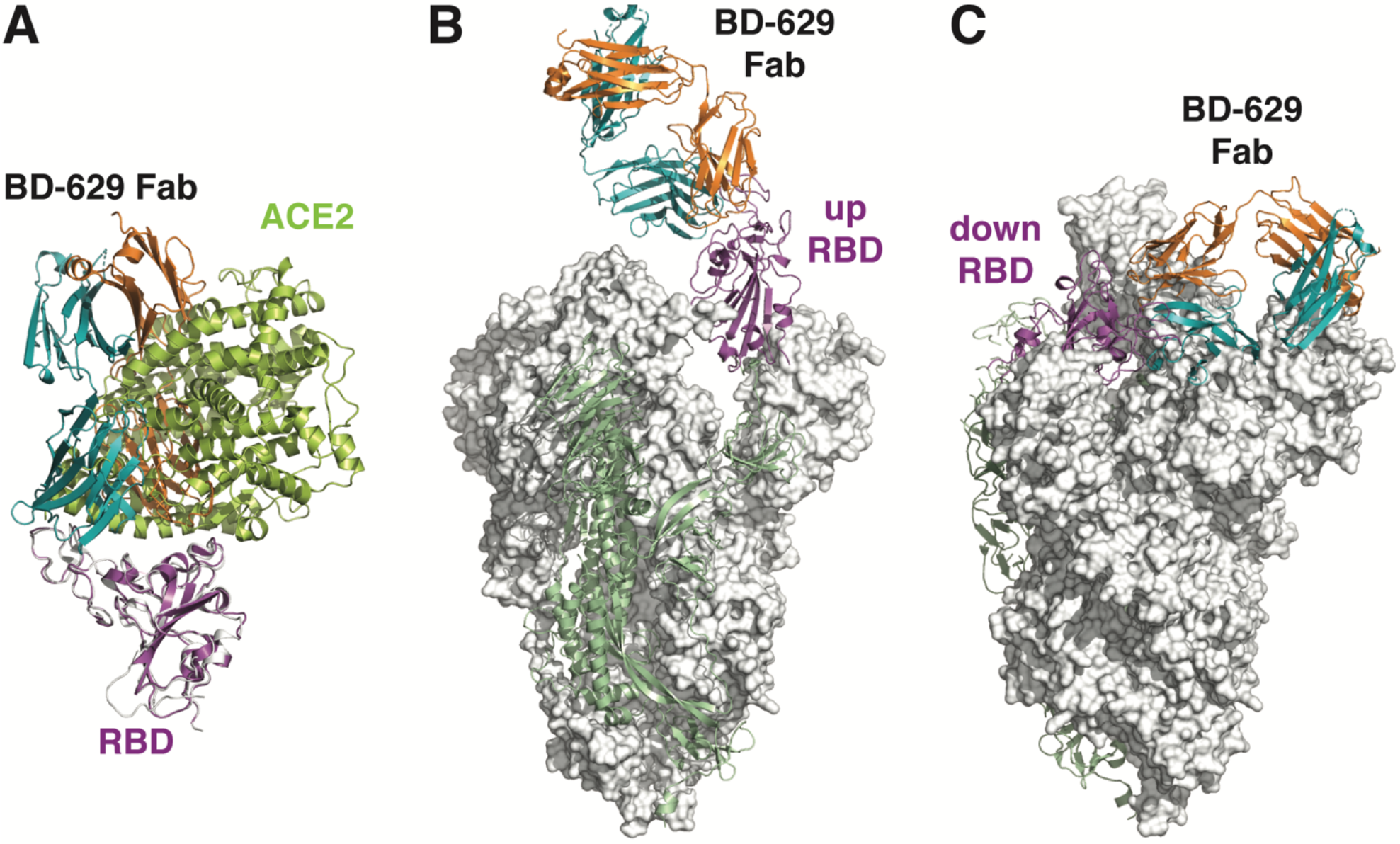
The *VH3-53/VH3-66* antibodies can only interact with the RBDs in the “up” conformation. (A) The *VH3-53/VH3-66* antibodies would block the interaction between RBD and ACE2. The structure of RBD/BD-629 Fab complex is overlaid onto that of the RBD/ACE2 complex (PDB ID: 6LZG), revealing that the epitope of BD-629 largely overlaps with the binding site of ACE2. RBD and ACE2 in the RBD/ACE2 complex are shown in white and lemon, respectively. The RBD/BD-629 Fab complex is shown using the same colors as in Figure 2. (B) BD-629 Fab is modeled onto the “up” RBD in the S trimer (PDB ID: 6VSB) by structural superimpositions. The “up” RBD is shown in magenta, and the rest of that S protomer is shown in green. The other two S protomers are shown on white surfaces. (C) BD-629 Fab is modeled onto one of the “down” RBD in the same S trimer structure. The “down” RBD is not available for the interaction with BD-629, due to the steric hindrance imposed by an adjacent protomer.

### BD-368-2’s epitope does not overlap with convergent antibodies’ binding site

Among the neutralizing antibodies we identified, BD-368-2 is the most potent, exhibiting an IC_50_ of 1.2 and 15 ng/mL against pseudo and authentic SARS-CoV-2 (Cao et al., 2020). BD-368-2 also showed high therapeutic and prophylactic efficacy in SARS-CoV-2-infected mice. The immunoglobulin heavy chain is encoded by germline genes *IGHV3-23*, *IGHD3-16*, and *IGHJ4*, respectively; whereas the light chain is encoded by *IGKV2-28* and *IGKJ5*. To characterize the molecular interactions between BD-368-2 and RBD, we first attempted to obtain the crystal structure of BD-368-2 Fab in complex with RBD. However, this endeavor was unsuccessful despite extensive trials. We then discovered that BD-368-2 Fab could bind to RBD together with the Fabs of several *VH3-53/VH3-66* antibodies. We subsequently determined the crystal structures of three tripartite complexes consisting of the Fabs of these antibodies and RBD: BD-236/RBD/BD-368-2, BD-604/RBD/BD-368-2, and BD-629/RBD/BD-368-2. The resolutions are 3.4 Å, 2.7 Å, and 2.7 Å, respectively (Table S2).

These tripartite complexes display Y-like shapes, with the BD-368-2 Fab and *VH3-53/VH3-66* Fab attacking the RBD from opposite sides (Figure 4; Figure S5A-B). The interactions between the *VH3-53/VH3-66* Fabs and RBD are highly similar to those seen in the binary complexes described above (Figure S5C-E). Five regions in the BD-368-2 Fab are involved in interacting with RBD: heavy chain CDRH1 and CDRH3, DE loop in the VH domain, and light chain CDRL1 and CDRL2 (Figure 5A). The remaining two CDRs, especially CDRH2, do not directly contact RBD, suggesting that the interaction between BD-368-2 and RBD could be further enhanced by structure-based protein engineering. Gly26, Phe27, and Ala28 in CDRH1 cradle Tyr449^S^ (Figure 5B). Tyr32 in CDRH1 and Arg102 in CDRH3 together form robust packing with Phe490^S^. Arg102 also attaches to Glu484^S^ via a bidentate interaction. In addition, several hydrogen bonds are present between the heavy chain CDRs and RBD, involving heavy chain residues Arg100, Tyr105, Asp106, and RBD residues Gly482^S^, Glu484^S^ (Figure 5B). Ser75 and Asn77 in the DE loop of the VH domain form hydrogen bonds with Arg346^S^ and Asn450^S^ (Figure 5C). The light chain of BD-368-2 Fab mainly plays a supportive role in stabilizing the conformation of the heavy chain residues. Direct interactions between the light chain and RBD are seen between Asn33 in CDRL1 and Asn481^S^, which form reciprocal hydrogen bonds between their respective main chain and side chain groups (Figure 5D). Asn33, Tyr35, Tyr37, and Leu55 together create a pocket to accommodate Val483^S^.

**Figure 4.**
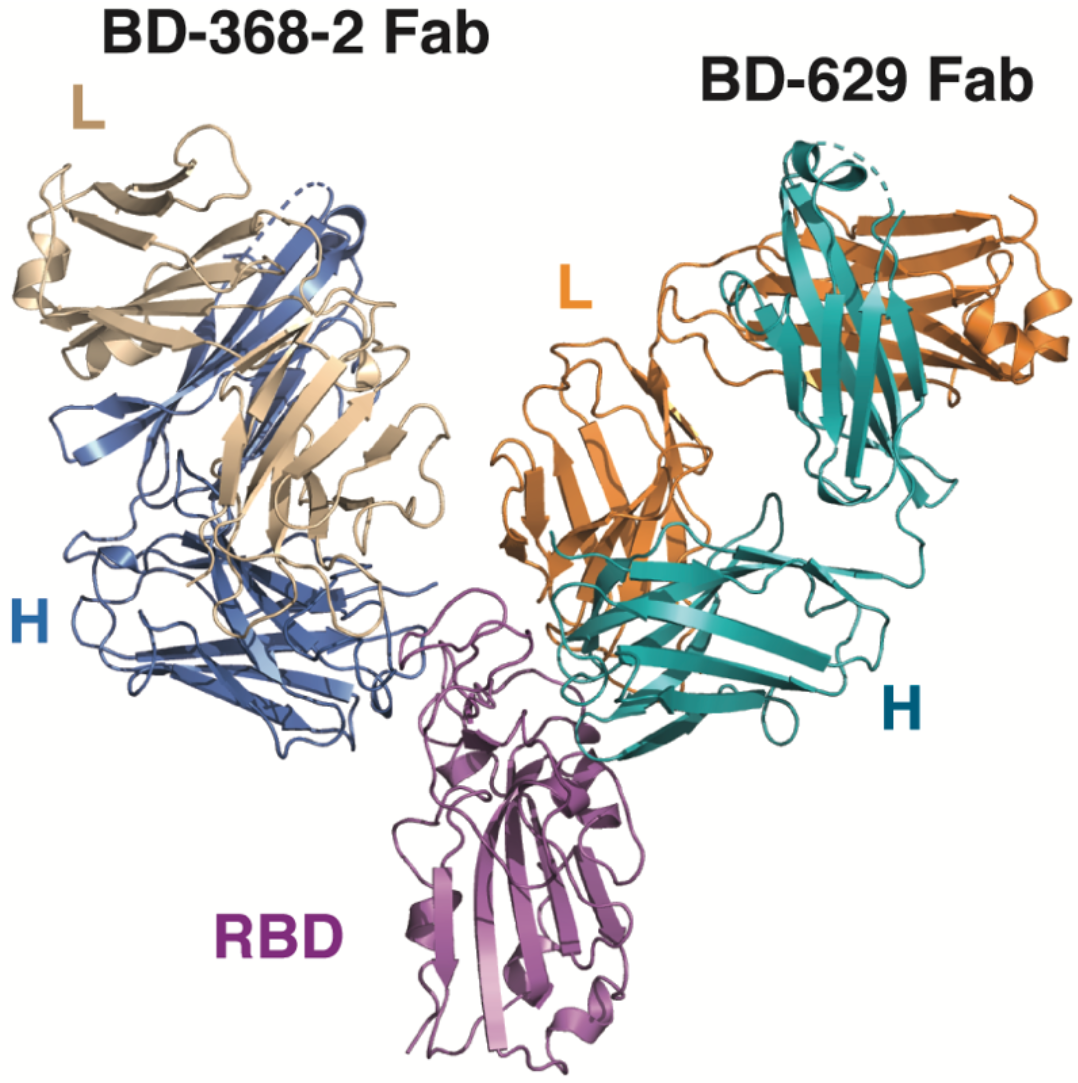
BD-368-2 can bind to the RBD together with the *VH3-53/VH3-66* antibodies. The crystal structure of RBD in complex with the Fabs of both BD-368-2 and BD-629 is shown in ribbon diagrams. The heavy chain and light chain of BD-368-2 Fab are shown in marine and wheat, respectively. RBD and BD-629 Fab are shown using the same color scheme as in Figure 2.

**Figure 5.**
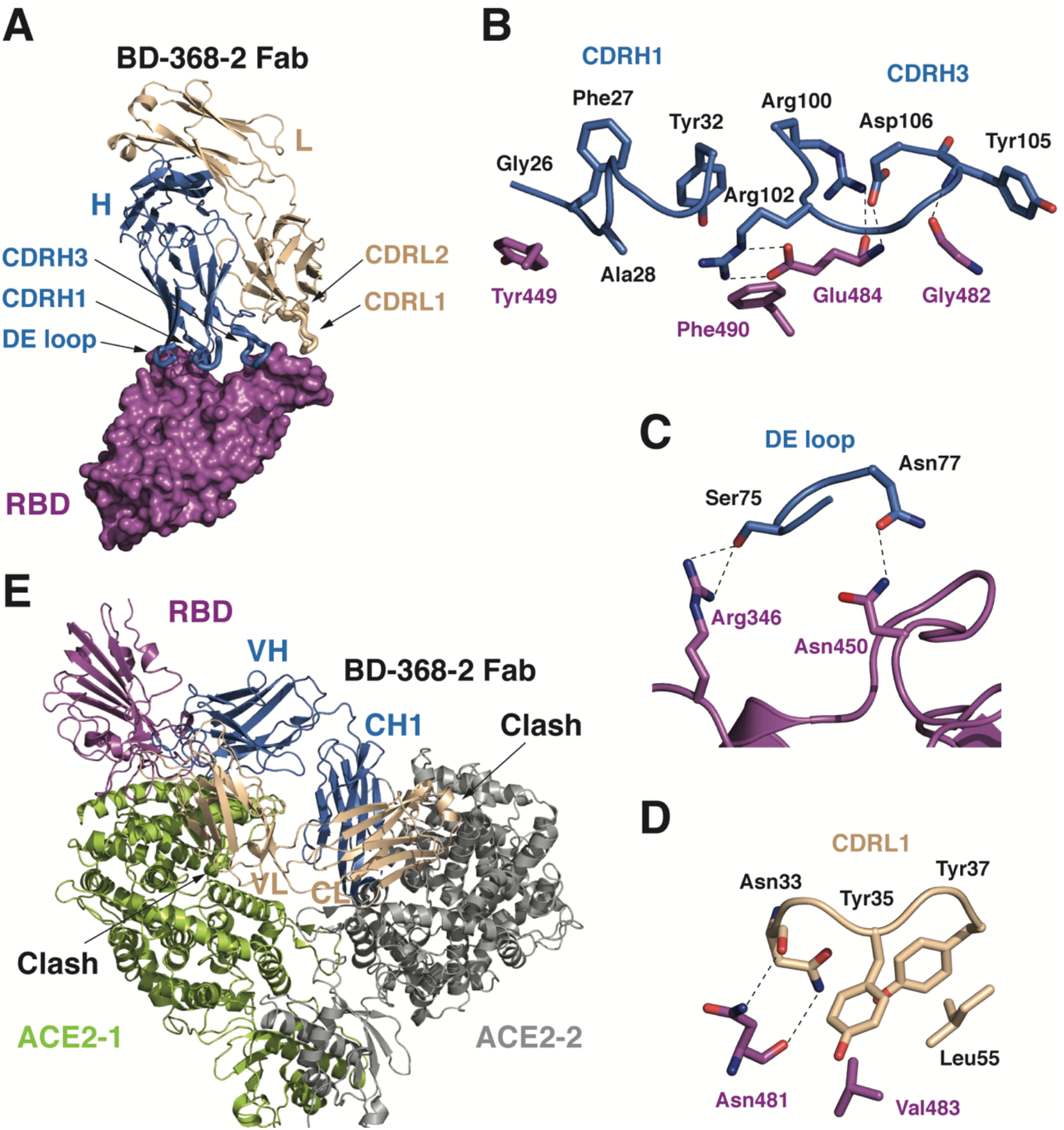
The interactions between BD-368-2 Fab and RBD. (A) RBD is shown in a surface view. BD-368-2 Fab is shown in ribbons. The five regions in BD-368-2 that interact with RBD are highlighted using thicker ribbons. (B) Interactions between CDRH1, CDRH3, and RBD. Polar interactions are indicated by dashed lines. (C) Interactions between the DE loop in the BD-368-2 VH domain and RBD. (D) Interactions between the BD-368-2’s VL domain and RBD. (E) BD-368-2 impedes the interaction between RBD and ACE2. The structure of RBD in the RBD/BD-368-2 complex is overlaid onto the RBD in the RBD/ACE2/B^0^AT1 complex (PDB ID: 6M17). BD-368-2 would clash with both protomers in the ACE2 dimer and therefore interfere with the interaction between RBD and ACE2.

The epitope of BD-368-2 does not significantly overlap with the binding site of ACE2 on RBD. Nevertheless, a structural superimposition of the RBD/BD-368-2 and RBD/ACE2 complexes reveals a clash between the VL domain of BD-368-2 Fab and ACE2 (Figure 5E), consistent with our previous analyses that BD-368-2 competitively inhibits the interaction between RBD and ACE2 (Cao et al., 2020). Furthermore, ACE2 exists as a homodimer in vivo (Yan et al., 2020). The constant domains in the BD-368-2 Fab would significantly clash with the other ACE2 protomer in the ACE2 dimer as well. The presence of the Fc region in RBD-BD-368-2 IgG would cause even more pronounced steric obstruction. Thus, BD-368-2 can directly block the interaction between RBD and ACE2, thereby exerting a protective effect.

### Cryo-EM structure of BD-368-2 in complex with the prefusion-stabilized S trimer

To further investigate the molecular mechanism by which BD-368-2 neutralizes SARS-CoV-2, we set to characterizing its interaction with the S trimer using cryo-EM. In the beginning, we used the 2P variant of the S protein (S-2P), which was designed by McLellan’s group and contains two stabilizing proline substitutions at residues 986-987 (Wrapp et al., 2020). We have successfully used this mutant to determine the cryo-EM structure of the S trimer in complex with the Fab of BD-23 (Cao et al., 2020). However, BD-368-2 Fab promptly disrupted the prefusion state structure of S-2P (Figure 6A). This phenomenon is reminiscent of S230, an antibody isolated from a person infected by SARS-CoV, which can functionally mimic ACE2 and promote a fusogenic-like conformational rearrangement of SARS-CoV S (Walls et al., 2019).

**Figure 6.**
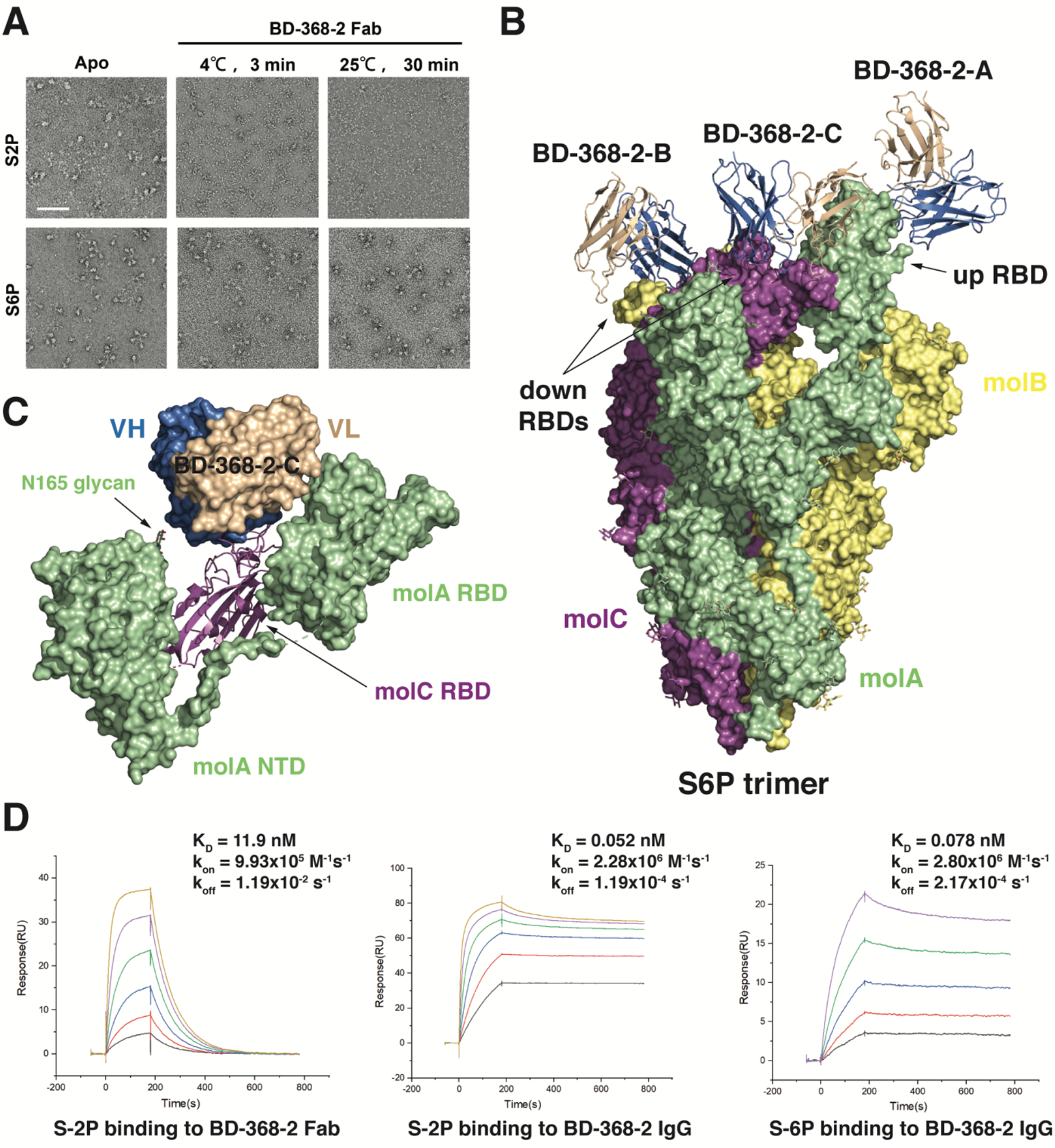
Cryo-EM structure of BD-368-2 Fabs in complex with the S-6P trimer. (A) BD-368-2 Fab induced significant structural changes of S-2P, as assessed by the negative stain EM. S-2P and S-6P both exist as stable trimers by themselves. Upon the treatment of BD-368-2 Fab, S-2P exhibits significant structural changes, whereas S-6P is more stable. The scale bar is 100 nm. (B) Cryo-EM structure of the S-6P trimer in complex with three BD-368-2 Fabs reconstructed at 3.5 Å. The S-6P trimer is depicted using a surface representation with the three protomers shown in green (molA), yellow (molB), and magenta (molC), respectively. The RBD in molA exhibits an “up” conformation, whereas RBDs in molB and molC are “down”. The Fv region of BD-368-2 Fab is shown in marine and wheat ribbons. (C) BD-368-2-C, which mainly interacts with the RBD in molC, appears to also contact the NTD and RBD in molA. (D) Surface plasmon resonance sensorgrams showing the binding between the S trimers and BD-368-2 Fab or IgG.

To this end, we produced S-HexaPro (S-6P), which was reported by the McLellan group recently (Hsieh et al., 2020). S-6P has a more stabilized prefusion state due to the introduction of four additional proline substitutions in the S2 segment of the S protein (F817P, A892P, A899P, A942P), which likely hinder the conformational change of S2. Indeed, S-6P is much more stable compared to S-2P in the presence of BD-368-2 Fab (Figure 6A). We subsequently determined a cryo-EM structure of BD-368-2 Fab in complex with S-6P at an overall resolution of 3.5 Å (Figure S6, Table S3).

S-6P exhibits an asymmetric conformation as previously observed (Cao et al., 2020; Wrapp et al., 2020), with one RBD “up” and two RBDs “down” (Figure 6B). All three RBDs are engaged by the BD-368-2 Fabs. Notably, one BD-368-2 Fab (BD-368-2-C, Figure 6C) that binds to one of the “down” RBD is spatially sandwiched between the NTD and RBD of the adjacent RBD-“up” protomer. The VH domain of this Fab is close to a glycan attached to Asn165 in the NTD of the RBD-“up” protomer, whereas the VL domain appears to contact the “up” RBD directly. In a way, it seems that besides its own RBD target, this Fab is also exploiting the adjacent S protomer to gain further avidity. Together, our structural analyses suggest that BD-368-2 can bind to the RBDs regardless of their “up” and “down” positions to achieve full occupancy of the S trimer. Consistently, BD-368-2 IgG exhibits markedly increased binding affinities for the S trimer compared to its Fab, likely because of the multivalent interactions (Figure 6D).

## Discussion

Here we performed a systematic structural analysis of the SARS-CoV-2 NAbs. Our results shed light on their neutralizing mechanisms. Both *VH3-53/VH3-66* NAbs and BD-368-2 directly prevent RBD from binding to the human receptor ACE2, thus fending the cells off the viral particles. In contrast to the *VH3-53/VH3-66* antibodies that can only engage the “up” RBD, BD-368-2 is unique in the way that it can access its epitopes on the S protein regardless of the “up” and “down” positions of the RBDs. We further show that BD-368-2 promptly disrupts the prefusion state of the S protein, which likely reflects a premature fusogenic-like structural rearrangement. This is highly evocative of the neutralizing mechanism proposed for S230 (Walls et al., 2019). A similar mechanism has been proposed recently for CR3022 (Huo et al., 2020), another neutralizing antibody isolated from a convalescent SARS patient that can cross-react with the S protein of SARS-CoV-2 (Tian et al., 2020; Yuan et al., 2020b).

We further show that the epitopes of the *VH3-53/VH3-66* NAbs and BD-368-2 have no overlaps, and can engage one RBD simultaneously. These results provide a foundation for combination therapy. In fact, BD-368-2 may further potentiate the activity of the *VH3-53/66* antibodies, since it can induce rapid structural changes of the S trimer, which may lead to the exposure of the RBDs that were originally in the “down” state. The simultaneous use of two antibodies targeting different epitopes of the S protein can not only potentially lead to more effective treatments, but also prevent the emerging of mutant viruses that escape from the neutralizing power of one antibody. Recently, scientists at Regeneron have described such a pair of antibodies, REGN10987 and REGN10933, and showed that their cocktail indeed prevented the generation of escaping mutants using a pseudovirus system (Baum et al., 2020; Hansen et al., 2020). Structural comparisons suggest that REGN10933 and the *VH3-53/VH3-66* antibodies such as BD-629 bind to a largely similar area on RBD, whereas REGN10987 and BD-368-2 each aims at a different region (Figure S7). BD-629 can bind to RBD together with REGN10987, whereas BD-368-2 would clash with both REGN10987 and REGN10933 (Figure S7C) and therefore can’t function in a pair with any of them.

Besides the *VH3-53/VH3-66* antibodies, further structural analyses suggest that BD-368-2 appears to be able to bind RBD together with two other antibodies: CR3022 and S309 (Figure 7). Like CR3022, S309 was also originally isolated from a convalescent SARS patient and can neutralize SARS-CoV-2 (Pinto et al., 2020).

**Figure 7.**
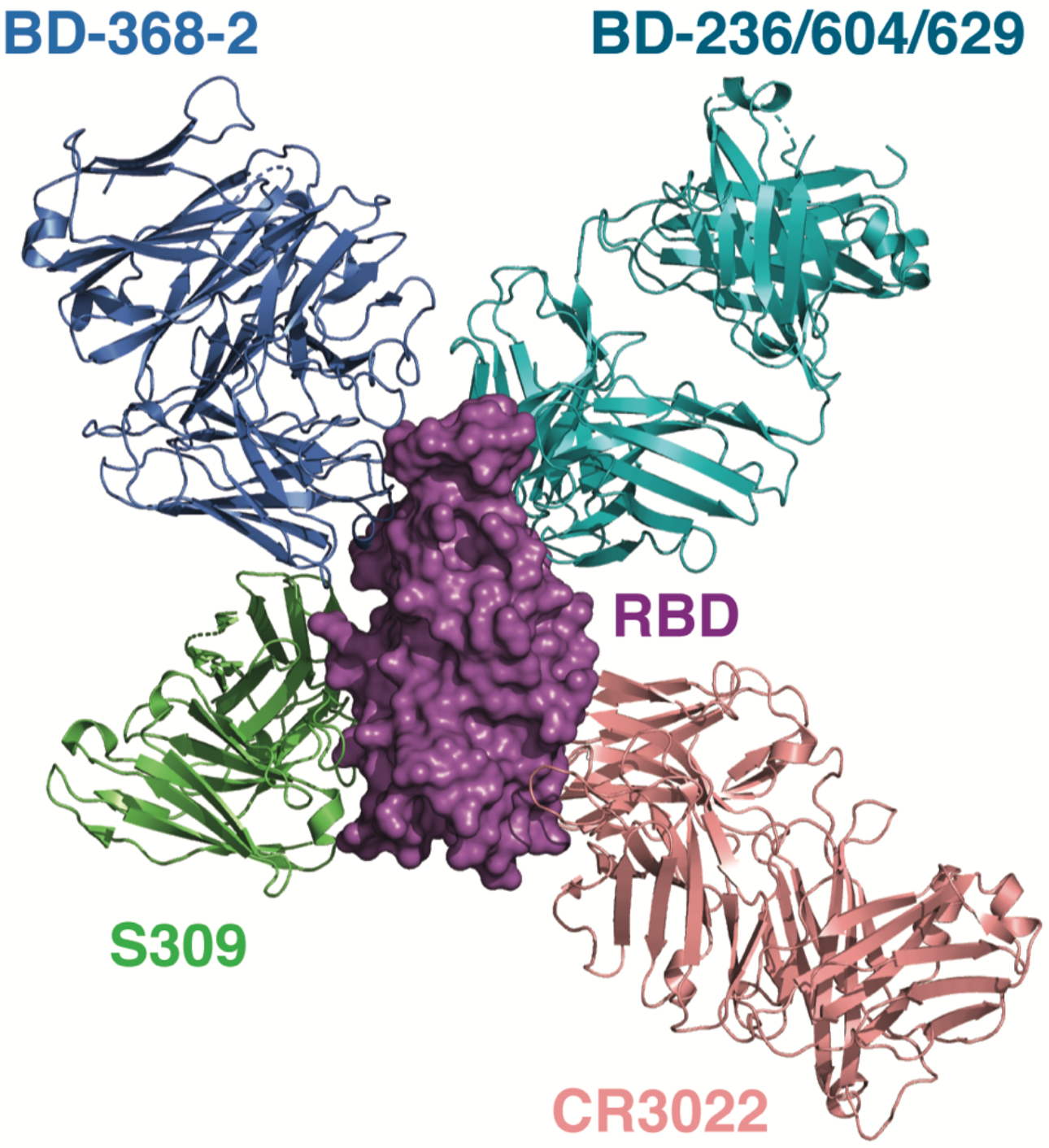
BD-368-2 can bind to the RBD together with the *VH3-53/VH3-66* antibodies, S309, and CR3022. The structures of the SARS-CoV-2 S in complex with S309 (PDB ID: 6WPS) and RBD in complex with CR3022 (PDB ID: 6W41) are superimposed onto the structure of BD-368-2/RBD/BD-629 to illustrate their binding modes. These antibodies have non-overlapping epitopes on RBD.

These antibodies each have a unique epitope and displays a distinct binding pose. Among them, BD-368-2 binds to the RBD regardless of its “up” and “down” state, blocks the engagement of ACE2, and causes drastic conformational changes of the S trimer. All these effects likely contribute to its potent neutralizing activity. S309 recognizes a glycan-containing epitope and can also bind both the “up” and “down” RBDs (Pinto et al., 2020). Nevertheless, S309 does not directly compete with ACE2 for binding to the S protein, and likely has a different mechanism of neutralization. The *VH3-53/VH3-66* antibodies engage the “up” RBDs to prevent their interaction with ACE2. CR3022 recognizes an epitope that is inaccessible in the prefusion state of the S protein, and have to engage the RBDs when at least two RBDs are “up” and also rotated (Huo et al., 2020; Yuan et al., 2020b). Certainly, the SARS-CoV-2 S protein is flexible in nature and exists in multiple conformations, and the presence of some of these antibodies can change the conformation landscape and trigger conformational changes, as shown for BD-368-2 (Figure 6A) and CR3022 (Huo et al., 2020). Collectively, these very different and non-overlapping antibodies provide an arsenal of therapeutic choices, and multiple combinations between them, even with antibodies that target the NTD of the S protein (Chi et al., 2020; Liu et al., 2020), can be tested to intervene the SARS-CoV-2 pandemic.

## KEY RESOURCES TABLE

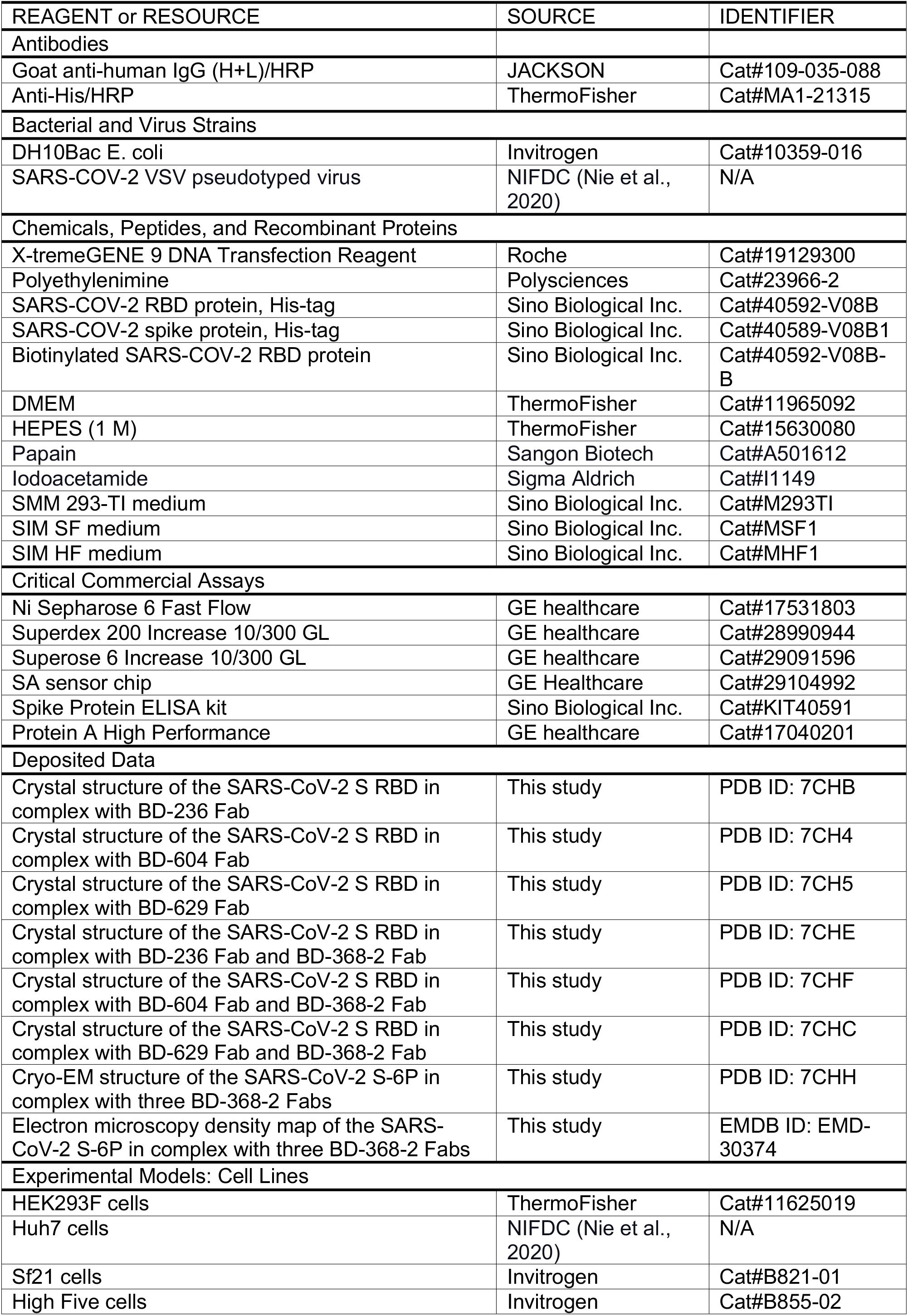
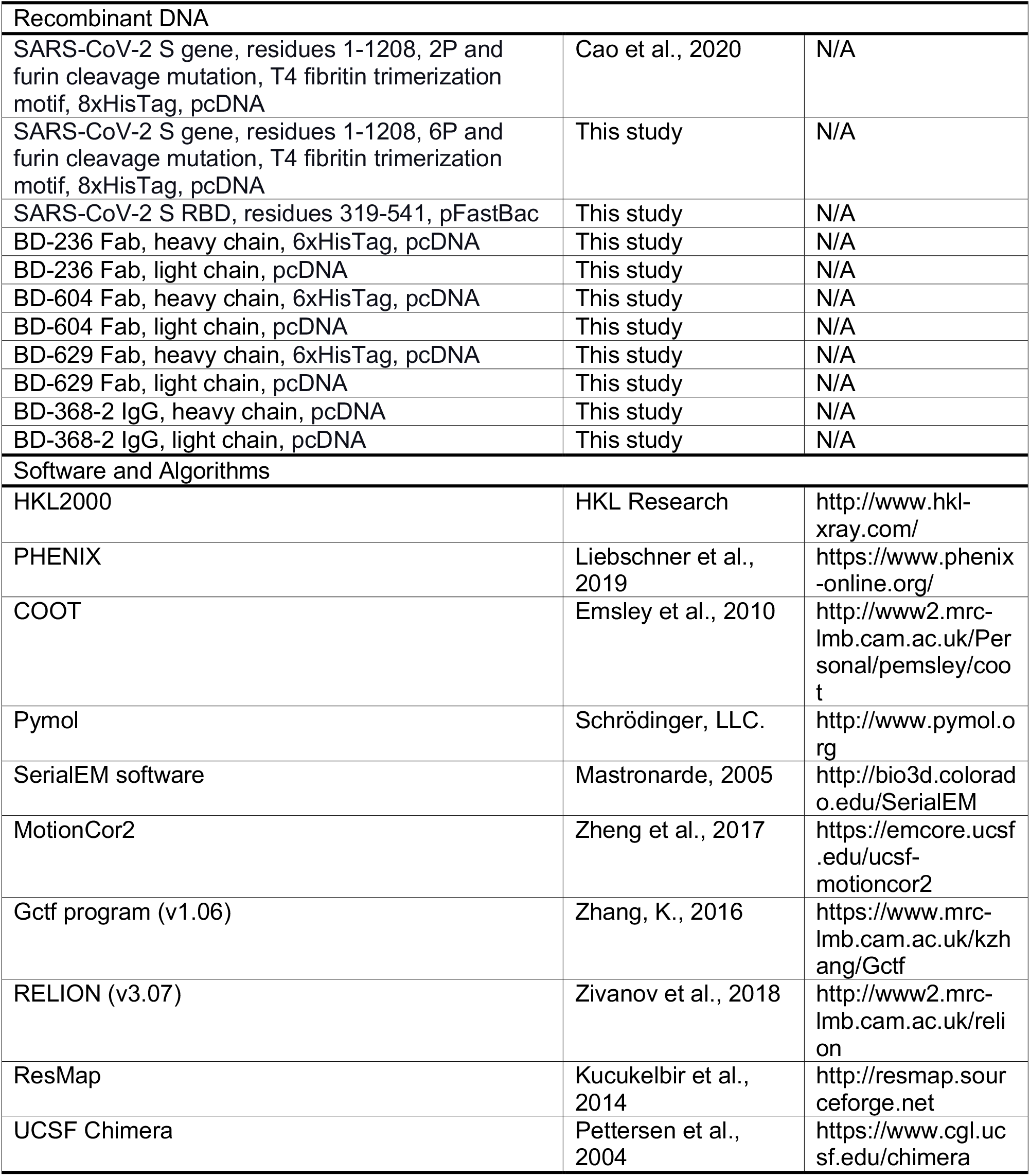

## RESOURCE AVAILABILITY

### Lead Contact

Further information and requests for resources and reagents should be directed to and will be fulfilled by the Lead Contact, Junyu Xiao (J.X.)

### Material Availability

There are restrictions on the availability of antibodies due to limited stock and continued consumption. We are glad to share remaining antibodies with reasonable compensation for processing and shipping upon completion of a Material/Data Transfer Agreement for non-commercial usage.

### Data and Code Availability

Human antibody sequences are available on the European Genome-Phenome Archive upon publication. Material/Data Transfer Agreements, which allow the use of the antibody sequences for non-commercial purposes but not their disclosure to third parties, are needed to obtain the sequences by contacting the Data Access Committee.

## STAR Methods

### *In Vitro* expression of the antibodies and ELISA quantification

All antibody sequences in this manuscript were generated in the previous study (Cao et al., 2020). The antibodies were *in vitro* expressed using HEK293 cells, and the binding specificities were quantity by ELISA against the Spike protein and the RBD protein, as described previously. An antibody is defined as ELISA-positive when the OD_450_ is saturated using 1 μg/mL RBD/S protein.

### Surface plasmon resonance

The dissociation coefficients for the binding between BD-368-2 and the S trimers were measured using a Biacore T200 (GE Healthcare) as previously described (Cao et al., 2020). Anti-His-tag antibodies were loaded to a SA sensor chip by NHS to capture the His-tag labeled S trimer. Serial dilutions of purified BD-368-2 Fab or IgG were then injected, ranging in concentrations from 50 to 0.78 nM. The running buffer is phosphate buffered saline, pH 7.4, supplemented with 0.005% (v/v) P20. The resulting data were fit to a 1:1 binding model using the Biacore Evaluation Software.

### Measurement of antibody neutralization potency

The pseudovirus neutralization assays were performed using Huh-7 cell lines, as described previously (Cao et al., 2020). Briefly, various concentrations of antibodies (3-fold serial dilution using DMEM) were mixed with the same volume of SARS-CoV-2 pseudovirus in a 96 well-plate. The mixture was incubated for 1 h at 37 °C and supplied with 5% CO_2_. Pre-mixed Huh-7 cells were added to all wells and incubated for 24 h at 37 °C and supplied with 5% CO_2_. After incubation, the supernatants were removed, and D-luciferin reagent (Invitrogen) was added to each well and measured luciferase activity using a microplate spectrophotometer (Perkinelmer EnSight). The inhibition rate is calculated by comparing the OD value to the negative and positive control wells. IC_50_ and IC_80_ were determined by a four-parameter logistic regression using GraphPad Prism 8.0 (GraphPad Software Inc.).

### Protein expression and purification

The SARS-CoV-2 RBD (residues 319-541) with an N-terminal His_6_ tag was cloned into a modified pFastBac vector (Invitrogen) that encodes a gp67 signal peptide. Bacmids were generated using the Bac-to-Bac system (Invitrogen). Baculoviruses were generated and amplified using the Sf21 insect cells. For protein production, Hi5 insect cells at 1.5 million cells/mL were infected with the RBD baculovirus. The conditioned media were harvested after 48 h, concentrated using a Hydrosart Ultrafilter, and exchanged into the binding buffer (25 mM Tris-HCl, pH 8.0, 200 mM NaCl). The RBD protein was purified first using the Ni-NTA resin (GE Life Sciences) and then a Superdex 200 Increase 10/300 gel filtration column (GE Life Sciences). The final buffer used for the gel filtration step contains 20 mM HEPES, pH 7.2, and 150 mM NaCl.

The Fabs of BD-236, BD-604, and BD-629 were obtained by transient transfection in HEK293F cells using polyethylenimine (Polysciences) when the cell density reached 1 million cells/mL. A C-terminal His_6_ tag was added to the heavy chains. Four days after transfection, the conditioned media were collected, and the Fabs were purified using the Ni-NTA resin and Superdex 200 Increase column similarly as the RBD. BD-368-2 IgG was expressed by transient transfection in HEK293F cells and purified from the conditioned media using a Protein A column (GE Life Sciences). To obtain the BD-368-2 Fab, BD-368-2 IgG (2 mg/mL) was digested with papain (0.1 mg/mL) for 2 hours at 37 °C, in a buffer containing 50 mM phosphate buffer saline, pH 7.0, 2 mM EDTA, and 5.5 mM cysteine. Digestion was quenched using 30 mM iodoacetamide at 30 °C for 30 min. The Fc region was removed by protein A chromatography, and the BD-368-2 Fab was further purified using the Superdex 200 Increase column and eluted using the final buffer.

The S-2P expression construct was previously described (Cao et al., 2020). The S-6P construct (Hsieh et al., 2020) was generated from S-2P using a PCR based method. The S-2P or S-6P plasmid was transfected into the HEK293F cells when the cell density reached 1 million cells/mL and expressed for four days. The S proteins were purified using the Ni-NTA resin, followed by a Superose 6 Increase 10/300 gel filtration column (GE Life Sciences), and eluted using the final buffer.

### Crystallization and structure determination

The BD-236/RBD, BD-604/RBD, BD-629/RBD, BD-236/RBD/BD-368-2, BD-604/RBD/BD-368-2, and BD-629/RBD/BD-368-2 complexes were obtained by mixing the corresponding protein components at equimolar ratios and incubated on ice for 2 hours. The assembled complexes were further purified using the Superdex 200 Increase column and eluted with the final buffer. Purified complexes were concentrated to 7-10 mg/ml for crystallization. The crystallization experiments were performed at 18 °C, using the sitting-drop vapor diffusion method. Diffraction-quality crystals were obtained in the following solution conditions:

BD-236/RBD: 0.1 M sodium citrate, pH 5.0, and 8% (w/v) polyethylene glycol 8000;
BD-604/RBD: 0.2 M potassium phosphate dibasic, pH 9.2, and 20% (w/v) polyethylene glycol 3,350;
BD-629/RBD: 0.1 M sodium citrate tribasic dihydrate, pH 5.0, and 18% (w/v) polyethylene glycol 20,000;
BD-236/RBD/BD-368-2: 0.1 M sodium acetate, pH 4.0, and 10% (w/v) polyethylene glycol monomethyl ether 2,000;
BD-604/RBD/BD-368-2: 0.2 M ammonium sulfate, 12% (w/v) polyethylene glycol 8000;
BD-629/RBD/BD-368-2: 0.1 M imidazole, pH 7.0, 20% (w/v) polyethylene glycol 6,000.

For data collection, the crystals were transferred to a solution containing the crystallization solution supplemented with 20% ethylene glycol or 20% glycerol before they were flash-cooled in liquid nitrogen. Diffraction data were collected at the Shanghai Synchrotron Radiation Facility (beamline BL17U) and the National Facility for Protein Science Shanghai (beamline BL19U). The data were processed using HKL2000 (HKL Research). All structures were solved by the molecular replacement method using the Phaser program (McCoy et al., 2007) in Phenix (Liebschner et al., 2019). The structural models were then manually adjusted in Coot (Emsley et al., 2010) and refined using Phenix.

### Negative staining electron microscopy

For the negative-staining study, S-2P, S-6P, and BD-368-2 Fab were diluted to 0.02 mg/ml using 25 mM HEPES, pH 7.2, 150 mM NaCl. BD-368-2 Fab was then mixed with S-2P or S-6P in a 1:1 volume ratio and incubated on ice for 3 min or at room temperature for 30 min. The mixture was then applied onto a glow-discharged carbon-coated copper grid (Zhong Jing Ke Yi, Beijing). After 1 min, the excess liquid was removed using a filter paper. The grids were then stained using 1% uranyl acetate for 30 seconds and air-dried. A Tecnai G2 20 Twin electron microscope (FEI) operated at 120 kV was used to examine the grids. Images were recorded using a CCD camera (Eagle, FEI).

### Cryo-EM data collection, processing, and structure building

Holy-carbon gold grids (Quantifoil, R1.2/1.3) were glow-discharged for 30 seconds using a Solarus 950 plasma cleaner (Gatan) with a 4:1 O_2_/H_2_ ratio. Four microliter S-6P (0.2 mg/mL) and 0.5 microliter BD-368-2 Fab (1.2 mg/mL) were mixed on ice for 1 minute, and then quickly applied onto the glow-discharged grids. Afterward, the girds were blotted with a filter paper (Whatman No. 1) at 4 °C and 100% humidity, and plunged into the liquid ethane using a Vitrobot Mark IV (FEI). The grids were first screened using a 200 kV Talos Arctica microscope equipped with a Ceta camera (FEI). Data collection was carried out using a Titan Krios electron microscope (FEI) operated at 300 kV. Movies were recorded on a K2 Summit direct electron detector (Gatan) using the SerialEM software (Mastronarde, 2005), in the super-resolution mode at a nominal magnification of 130,000, with an exposure rate of 7.1875 e-/Å^2^ per second. A GIF Quantum energy filter (Gatan) with a slit width of 20 eV was used at the end of the detector. The defocus range was set from – 0.7 to –1.2 μm. The micrographs were dose-fractioned into 32 frames with a total exposure time of 8.32 s and a total electron exposure of 60 electrons per Å^2^. Statistics for data collection are summarized in Table S3.

The workflow of data processing was illustrated in Figure S6. A total of 5,273 movie stacks were recorded. Raw movie frames were aligned and averaged into motion-corrected summed images with a pixel size of 1.055 Å by MotionCor2 (Zheng et al., 2017). The contrast transfer function (CTF) parameters of each motion-corrected image were estimated by the Gctf program (v1.06) (Zhang, 2016). Relion (v3.07) was used for all the following data processing (Zivanov et al., 2018). The S trimer (PDB ID: 6VSB) was used as a reference for the 3D classifications. The local resolution map was analyzed using ResMap (Kucukelbir et al., 2014) and displayed using UCSF Chimera (Pettersen et al., 2004).

The S trimer (PDB ID: 6VSB) and the Fv region of BD-368-2 Fab from the crystal structure described above were docked into the cryo-EM density using UCSF Chimera. Refinement was performed using the real-space refinement in Phenix. Figures were prepared using Pymol (Schrödinger) and UCSF Chimera.

## Acknowledgments

We thank the staff of the Shanghai Synchrotron Radiation Facility (beamline BL17U) and the National Facility for Protein Science Shanghai (beamline BL19U) for assistance with X-ray data collection; the Core Facilities at the School of Life Sciences, Peking University for help with negative-staining EM; the Cryo-EM Platform of Peking University for the assistance with EM data collection; the High-performance Computing Platform of Peking University for the assistance with computation. The work was supported by the National Key Research and Development Program of China (2017YFA0505200 to J.X.), the National Science Foundation of China (31822014 to J.X.), the Qidong-SLS Innovation Fund (to J.X.).

## Author contributions

X.S.X, X.D.S, and J.Y.X conceptualized the project, designed and coordinated the experiments. S.D and Q.Y.Z performed protein purification and crystallization experiments, with the help of X.X.D and H.X. Y.L.C led the NAbs discovery and characterization experiments. Q.Y.Z, H.X, B.W, C.G.J, and Q.S.W collected crystal diffraction data. Q.Y.Z and G.P.W prepared cryo-EM samples and collected data. G.P.W. processed the EM data, under the supervision of N.G. J.X. built the structural models and performed structural analyses. Y.L.C, X.S.X, X.D.S, and J.Y.X wrote the manuscript, with inputs from all the other authors.

---

**Figure S1.**
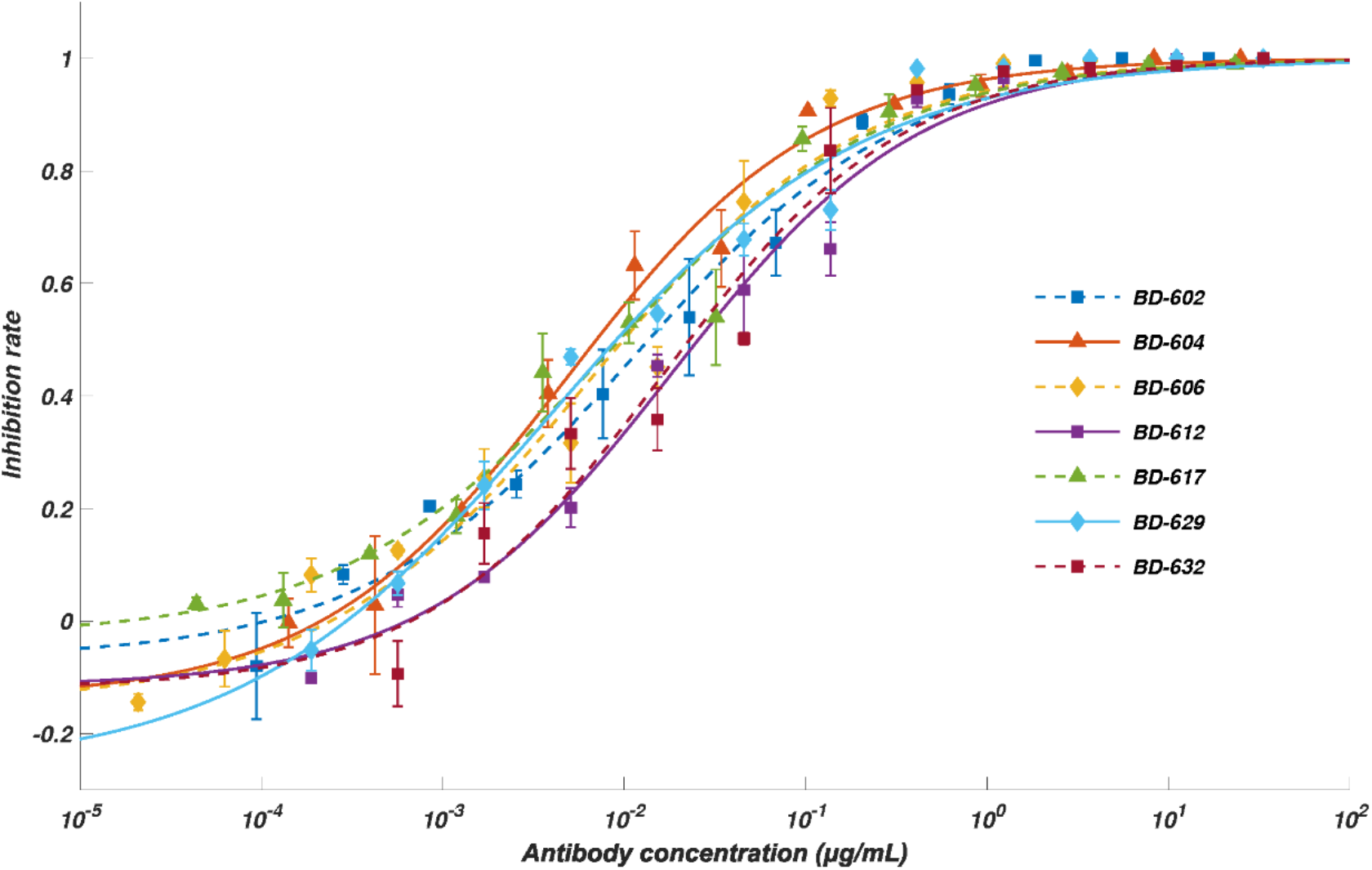
Neutralization ability of the potent *VH3-53/VH3-66* derived NAbs measured by SARS-CoV-2 pseudovirus. Related to Figure 1. Neutralization potency measured by a SARS-CoV-2 spike-pseudotyped VSV neutralization assay. Data for each NAb were obtained from a representative neutralization experiment, which contains three replicates. Data are represented as mean ± SD. IC_50_ and IC_80_ were calculated by fitting a four-parameter logistic curve.

**Figure S2.**
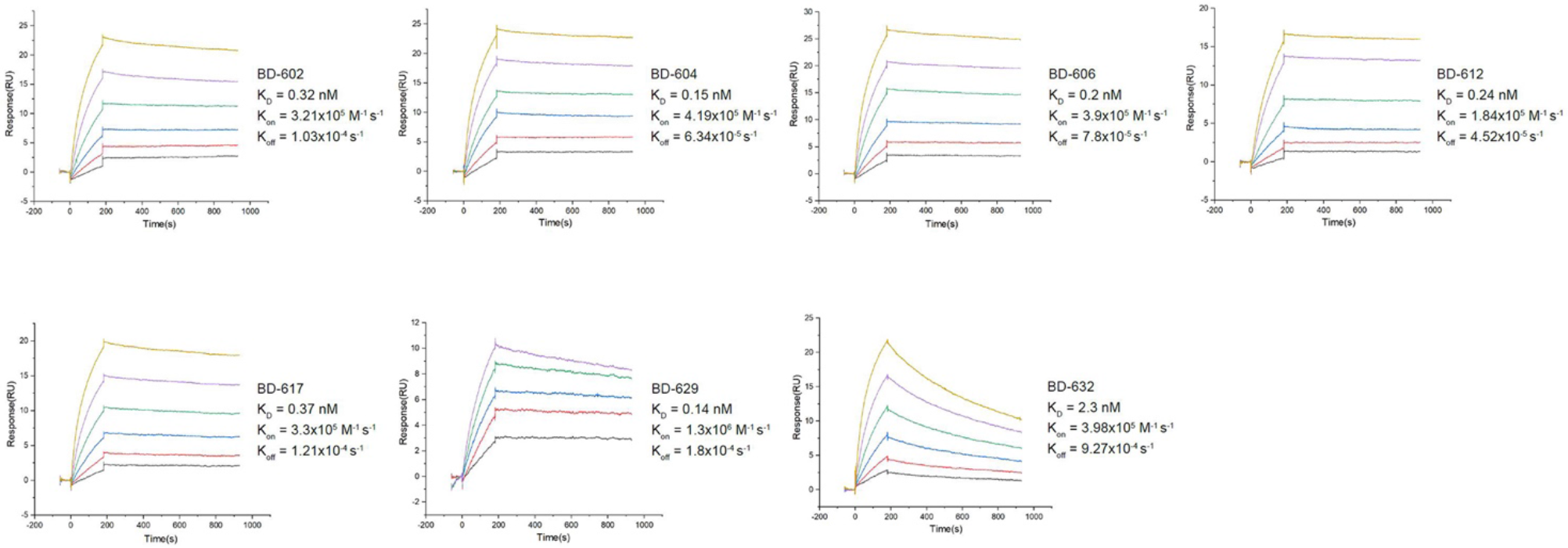
K_D_ measurement for the potent *VH3-53/VH3-66* derived NAbs. Related to Figure 1. Measurement of the dissociation constant against RBD for the representing NAbs. K_D_ is calculated using a 1:1 binding model. All analyses were performed by using a serial 2-fold dilution of biotinylated RBD, starting from 50 nM (yellow) to 1.56 nM (black).

**Figure S3.**
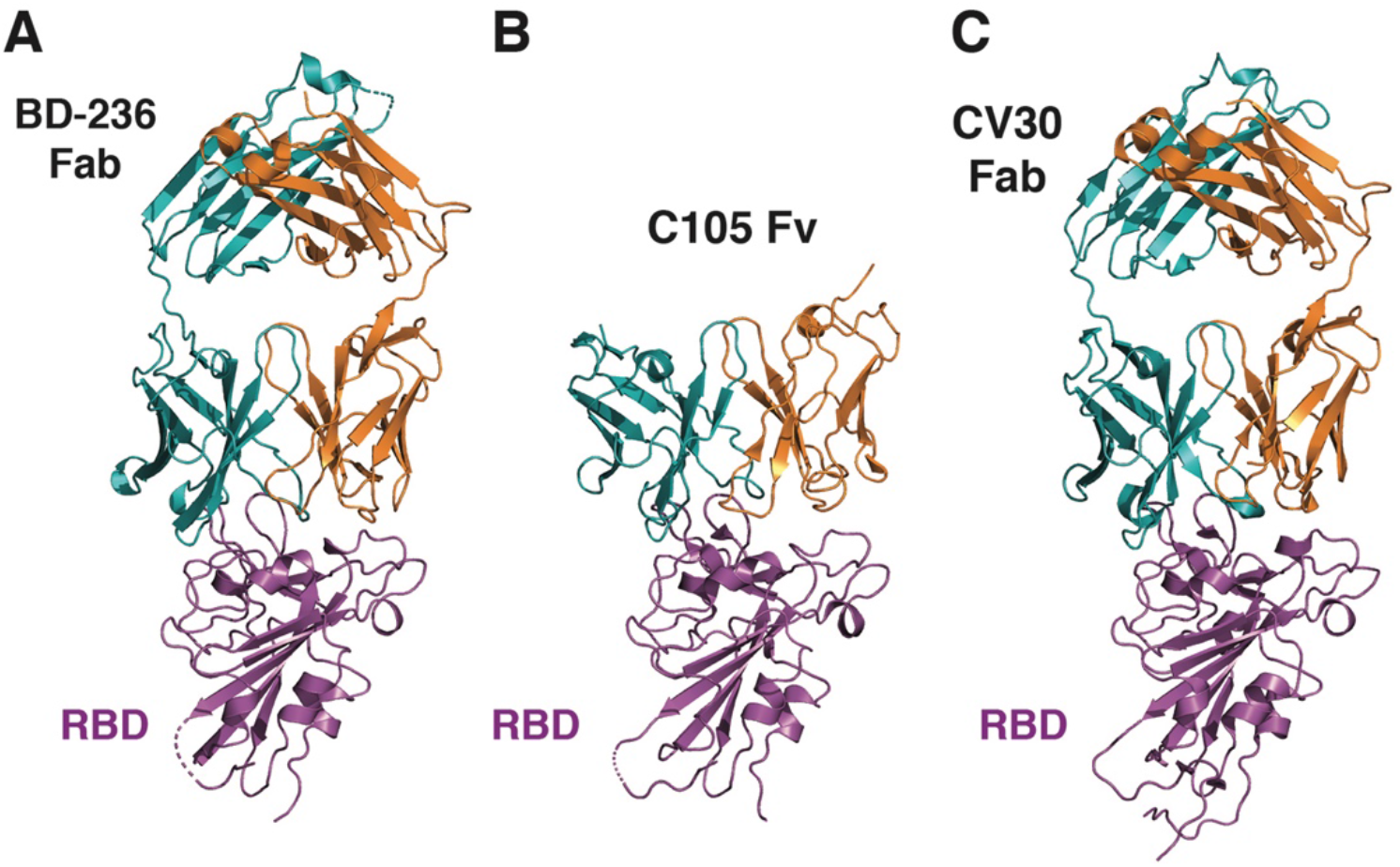
The *VH3-53/VH3-66* antibodies bind to RBD in a similar manner, Related to Figure 2. (A) The crystal structure of BD-236 Fab in complex with RBD. (B) C105 Fv in complex with RBD in the cryo-EM structure (PDB ID: 6XCM). (C) The crystal structure of CV30 Fab in complex with RBD (PDB ID: 6XE1).

**Figure S4.**
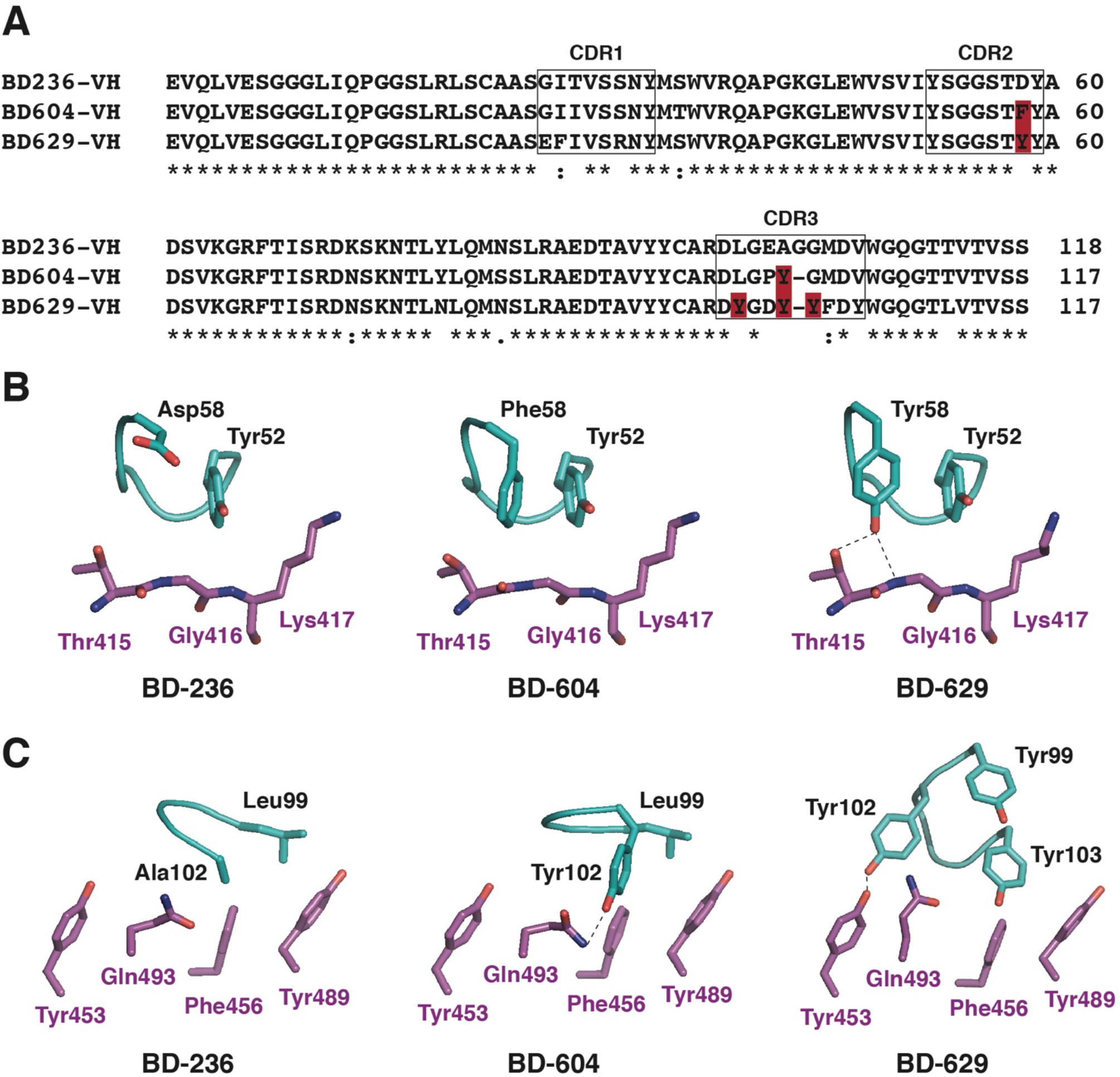
Structural comparison of the VH domains of the *VH3-53/VH3-66* antibodies, Related to Figure 2. (A) Sequence alignment of the VH domains of BD-236, BD-604, and BD-629. (B) Structure comparisons of the interactions with RBD mediated by the CDRH2s of these antibodies. (C) Structure comparisons of the interactions with RBD mediated by CDRH3s.

**Figure S5.**
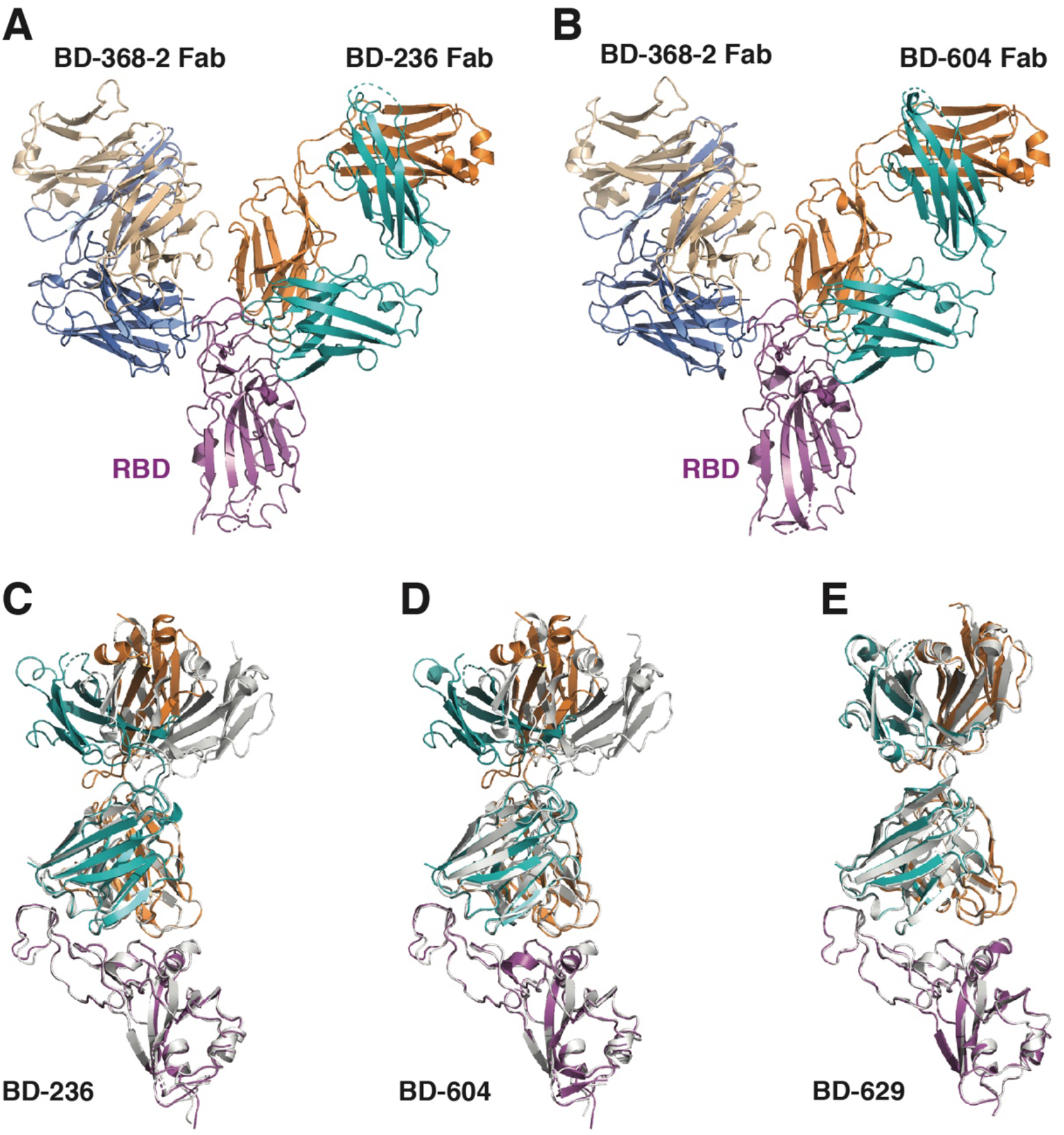
Structures of the *VH3-53/VH3-66* antibodies in binary and tripartite complexes with RBD and BD-368-2, Related to Figure 4. (A) Crystal structure of RBD in complex with the Fabs of both BD-368-2 and BD-236. (B) Crystal structure of RBD in complex with the Fabs of both BD-368-2 and BD-604. (C) Structural comparison of BD-236 in the BD-236/RBD binary and BD-236/RBD/BD-368-2 tripartite complexes. The binary complex structure, shown in white ribbons, is overlaid onto the tripartite complex shown in colored ribbons. BD-368-2 in the tripartite complex is not shown for clarity. RBD and the Fv region of BD-236 are largely superimposable in the two structures. A difference in the elbow angle of the BD-236 Fab is seen between the two structures due to flexibility. (D) BD-604 exhibits a similar structure in the tripartite and binary complexes except for the elbow angle. The binary complex is shown in white. (E) BD-629 exhibits a similar structure in the tripartite and binary complexes. The binary complex is shown in white.

**Figure S6.**
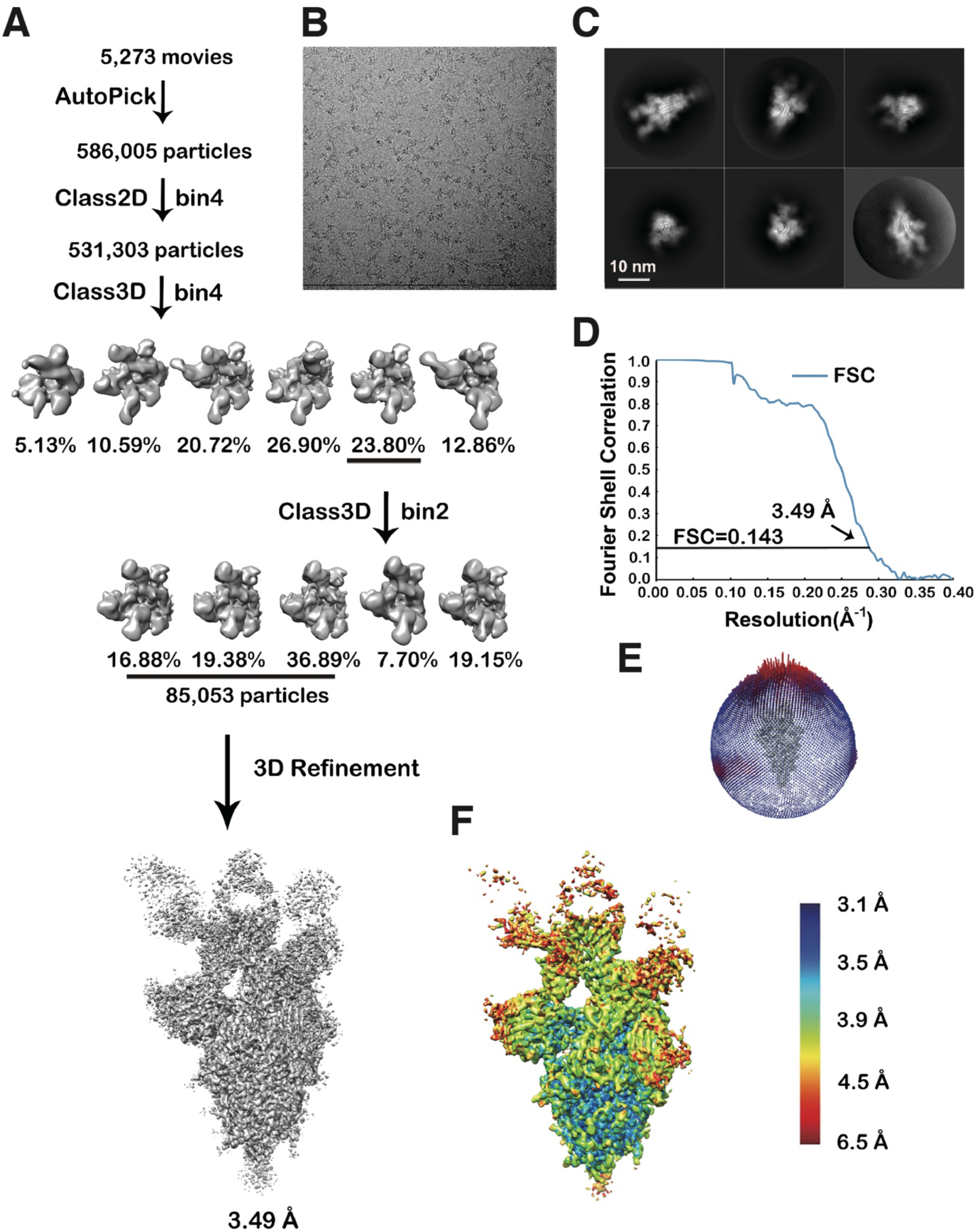
Workflow for the 3D reconstruction of the cryo-EM structure of the S-6P trimer in complex with three BD-368-2 Fabs, Related to Figure 6. (A) Flow chart of image processing. (B) A representative raw image collected using a Titan Krios 300 kV equipped with a K2 detector. (C) Representative 2D classes. (D) Gold standard Fourier shell correlation (FSC) curve with the estimated resolution. (E) Eulerian angle distribution of the particles used in the final 3D reconstruction. (F) Local resolution estimation of the final density map analyzed by ResMap.

**Figure S7.**
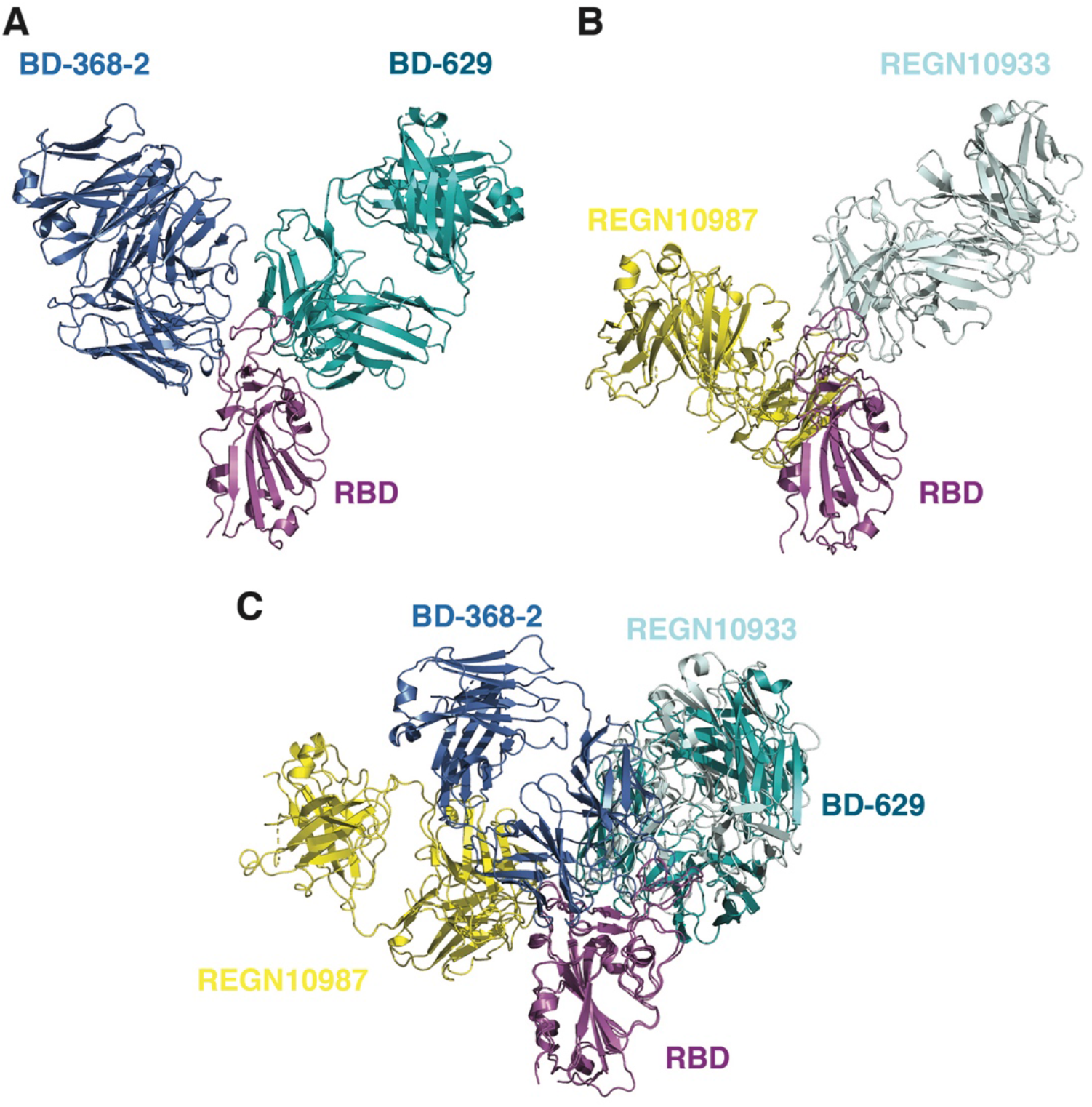
Structural comparisons of the BD-368-2/RBD/BD-629 and REGN10987/RBD/REGN10933 complexes, Related to Figure 7. (A) Crystal structure of the BD-368-2/RBD/BD-629 complex. BD-368-2, RBD, and BD-629 are shown in marine, magenta, and teal, respectively. (B) Cryo-EM structure of the REGN10987/RBD/REGN10933 complex (PDB ID: 6XDG). REGN10987, RBD, and REGN10933 are shown in yellow, magenta, and light blue, respectively. (C) Structural superimposition of the above two complexes.

